# Lokiarchaeon exhibits homoacetogenesis

**DOI:** 10.1101/826495

**Authors:** William D. Orsi, Aurèle Vuillemin, Paula Rodriguez, Ömer K. Coskun, Gonzalo V. Gomez-Saez, Gaute Lavik, Volker Morholz, Timothy G. Ferdelman

## Abstract

The proposed Asgard superphylum of Archaea comprises the closest archaeal relatives of eukaryotes, whose genomes hold clues pertaining to the nature host cell that acquired the mitochondrion at the origin of eukaryotes^1-4^. Genomes of the Asgard candidate Phylum ‘*Candidatus* Lokiarchaeota’ [Lokiarchaeon] suggest an acetogenic H_2_ –dependent lifestyle^5^ and mixotrophic capabilities^6^. However, data on the activity of Lokiarchaeon are currently lacking, and the ecology of the host cell that acquired the mitochondrion is debated^4,7^. Here, we show that in anoxic marine sediments underlying highly productive waters on the Namibian continental shelf Lokiarchaeon gene expression increases with depth below the seafloor, and was significantly different across a redox gradient spanning hypoxic to sulfidic conditions. Notably, Lokiarchaeon increased expression of genes involved in growth, carbohydrate metabolism, and the H_2_-dependent Wood-Ljungdahl (WLP) carbon fixation pathway under the most reducing (sulfidic) conditions. Quantitative stable isotope probing experiments revealed multiple populations of Lokiarchaeota utilizing both CO_2_ and diatomaceous extracellular polymeric substances (dEPS) as carbon sources over a 10-day incubation under anoxic conditions. This apparent anaerobic mixotrophic metabolism was consistent with the expression of Lokiarchaeon genes involved in transport and fermentation of sugars and amino acids. Remarkably, several Asgard populations were more enriched with ^13^C-dEPS compared to the community average, indicating a preference for dEPS as a growth substrate. The qSIP and gene expression data indicate a metabolism of “*Candidatus* Lokiarchaeota” similar to that of sugar fermenting homoacetogenic bacteria^8^, namely that Lokiarchaeon can couple fermentative H_2_ production from organic substrates with electron bifurcation and the autotrophic and H_2_-dependent WLP. Homoacetogenesis allows to access a wide range of substrates and relatively high ATP gain during acetogenic sugar fermentation^8^ thereby providing an ecological advantage for Lokiarchaeon in anoxic, energy limited settings.

---

The Benguela upwelling system is one of the most productive ecosystems of the world’s oceans and exhibits an oxygen minimum zone (OMZ) overlying the seafloor on the Namibian continental shelf^9^. Below the Namibian seafloor relatively high amounts of sulfide (H_2_S) and H_2_ are produced by sulfate reduction and microbial fermentation, respectively^10^. During the Southern Winter (July) of 2018 we sampled a 30 cm-long sediment core underlying hypoxic bottom waters of the Namibian OMZ (Extended Data Fig. 1). The sediments exhibited a redox gradient spanning hypoxic (ca. 25 µM O_2_) conditions at the seafloor surface to sulfidic conditions at 30 cm below seafloor (cmbsf) (Fig. 1a).

**Figure 1:**
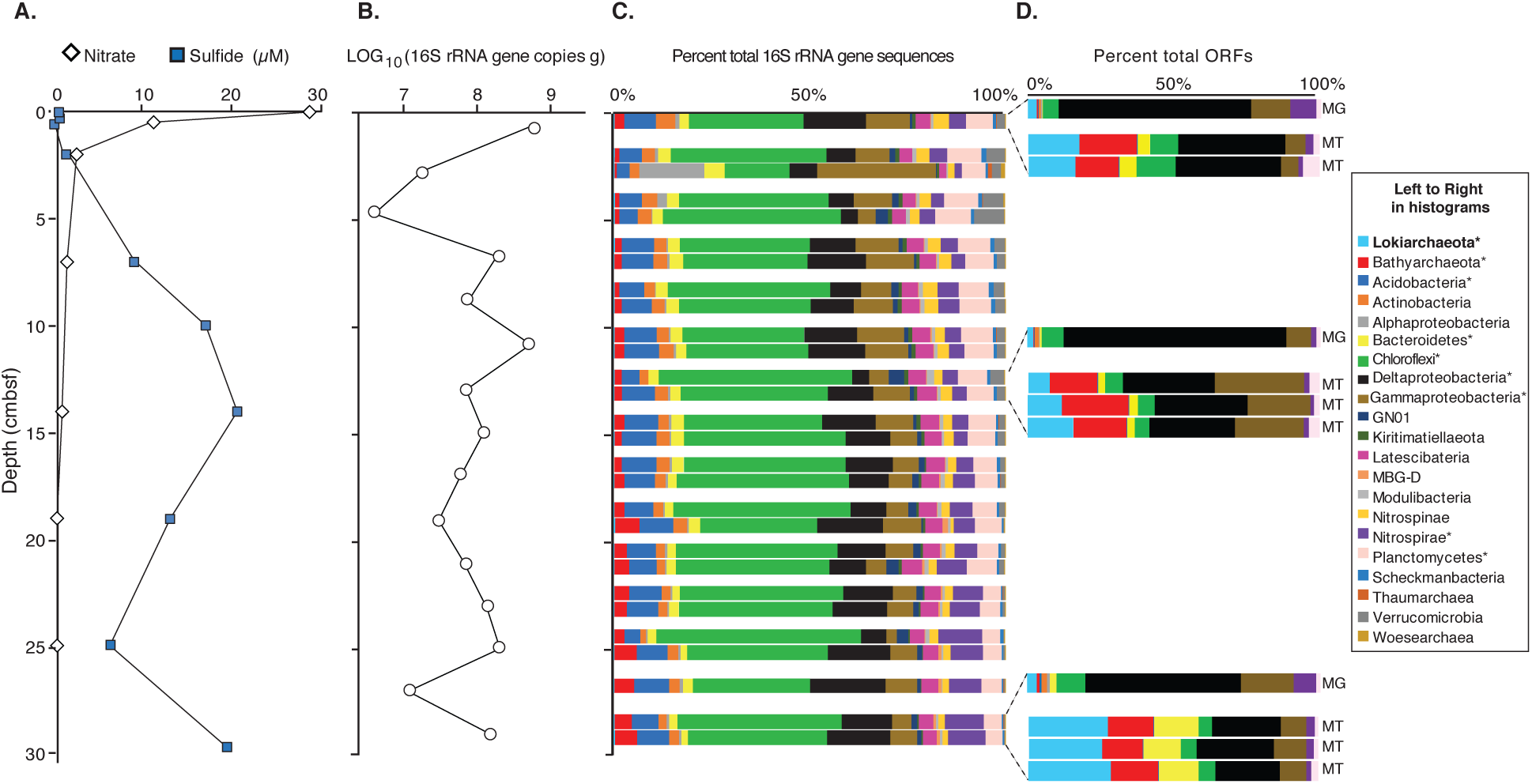
Lokiarchaeon gene expression increases in anoxic, sulfidic sediments. **(A)** Geochemical profile of nitrate (diamonds) and sulfide (squares) showing a redox transition zone between 8 and 12 cmbsf. Sulfide data represent average measurements from two cores. **(B)** Abundance of total 16S rRNA genes as determined by qPCR. **(C)** The relative abundance of 16S rRNA gene sequences (n=23,000 average per sample, SD: 3,000) assigned to microbial groups, closely spaced histograms are biological replicates. **(D)** Relative abundance of ORFs in replicate metatranscriptomes (MT; n=1,069 average per sample, SD: 238) with highest similarity to bacterial and archaeal groups. MG: Relative abundance of ORFs in unamplified metagenomes (n=57,526). Light blue = Lokiarchaeota. Asterisks indicate groups detected in 16S, metatranscriptome, and metagenome datasets.

Microbial abundance based on quantitative PCR (qPCR) of 16S rRNA genes showed that microbial biomass generally decreased with depth, with a subseafloor peak in biomass at the NO_3_^-^ - H_2_S interface. High-throughput sequencing of the 16S rRNA genes showed the presence of diverse microbial communities, dominated by the Chloroflexi (Extended Data Fig. S2) and Proteobacteria. In contrast, Archaea represented on average only 1.9% (SD: 0.6%, n=28 samples) of the 16S rRNA gene sequences (Fig. 1c). Phylogenetics of the 16S rRNA genes showed that the presence of Lokiarchaeota was attributed to two separate populations detected throughout the core (Extended Data Fig. S3). To check whether PCR primers were biasing the relative abundance of taxa, we performed direct (unamplified) sequencing of DNA extracted from three depths. Metagenomic analysis of these samples showed that protein encoding open reading frames (ORFs) assigned to Deltaproteobacteria were by far the most abundant at all depths, and that Archaea were 3.1% (SD: 1.1 %, n=3 samples) of ORFs (Fig 1d). The relative abundance of “*Candidatus* Lokiarchaeota” ORFs was much higher in the unamplified metagenomes (average 2.2%, SD=0.9, n=3 samples) compared to the relative abundance in 16S rRNA gene amplicon data (average%, SD: 0.004%, n=28 samples). The comparably lower abundance of Lokiarchaeon in the 16S rRNA gene sequences is likely due to PCR primer biases against the Asgard archaea. A relative abundance for “*Candidatus* Lokiarchaeota” (previously referred to as the “deep sea archaeal group”) at ca. 2% in our metagenomes is still relatively low compared to the abundance of this group in other studies of anoxic marine sediments where it can reach up to 10-50% relative abundance deeper in the subsurface^2,11^.

After applying metatranscriptomics to the same samples (Extended Data Table S1), expressed ORFs with highest similarity to predicted proteins from archaeal genomes were on average 36% (SD: 7%, n=8 samples) of total ORFs detected (Fig. 1d). Of these, on average 50% (SD: 11%, n=8 samples) had highest similarity to ‘*Candidatus* Lokiarchaeota’, which increased significantly (Welch Two Sample t-test: *P* = 0.01) up to 61% (SD: 0.2%, n=8 samples) in the deepest sample at 30 cmbsf (Fig. 1d). The relatively high amount of gene expression from Lokiarchaeon was inversely correlated with the abundance of Lokiarchaeon 16S and metagenomic datasets. We normalized the group-specific gene expression against the relative abundance of those same groups in the metagenomes (see Methods). This analysis showed that Archaea (Lokiarchaeon and Bathyarchaeota) had higher levels of gene expression compared to all other bacterial groups – with the exception of Bacteroidetes at 30 cmbsf. Moreover, Lokiarchaeon-specific gene expression was overexpressed at all depths relative to metagenomes, and increased with depth below the seafloor being greatest in the sulfidic interval at 30 cmbsf (Fig 2a). Genes with higher expression tended to have consistency between replicates, whereas those genes with lower expression have lower consistency between replicates, because the abundance of those transcripts are closer to our detection limit (Fig 2b). Moreover, across the entire redox spectrum sampled, gene expression of Lokiarchaeon was statistically significant (ANOSIM: *P* = 0.002, R=0.76) between the core top (hypoxic), 12 cmbsf (nitrate/sulfide interface), and 28 cm (sulfidic) (Fig. 2b). This pattern of gene expression is strongly suggestive of an anaerobic metabolism for Lokiarchaeon.

**Figure 2:**
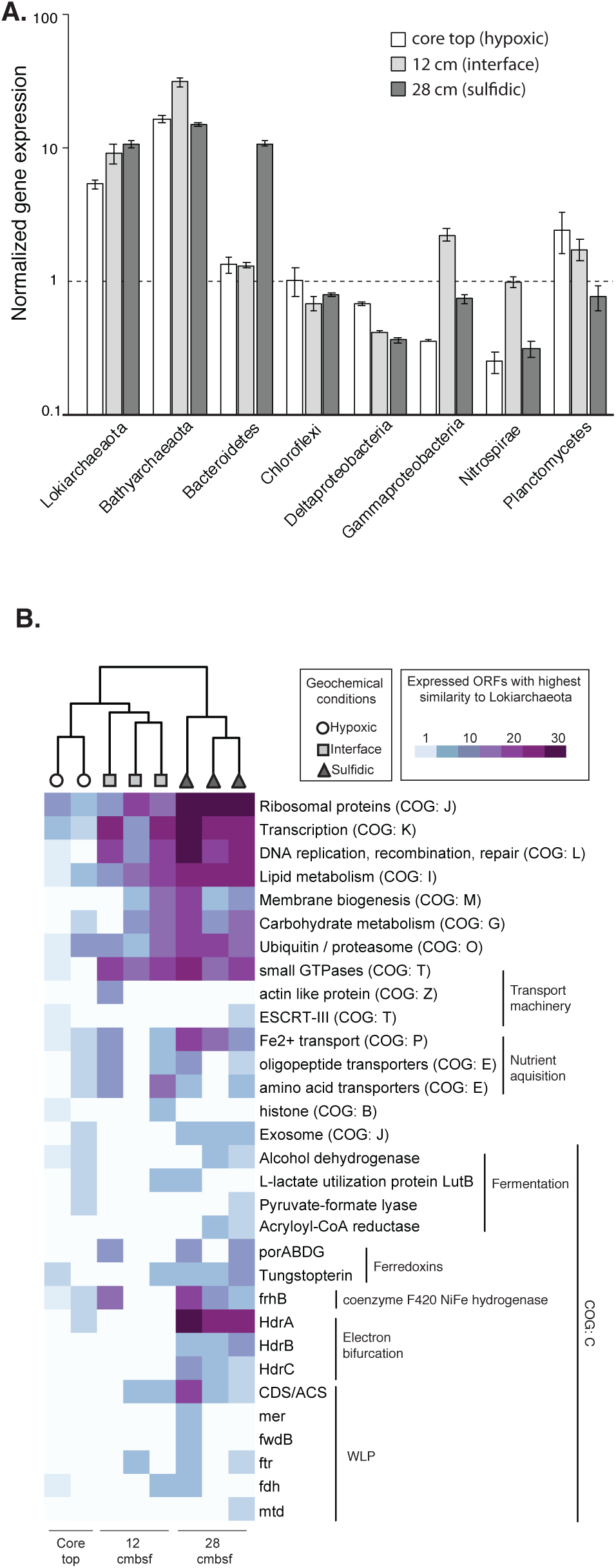
Lokiarchaeon gene expression across a hypoxic-sulfidic redox gradient. **(A)** The y axis represents gene expression normalized against group abundance: (% total expressed ORFs per group / % total ORFs per group in metagenomes). Error bars represent ranges in normalized values across biological replicates. Values above or below the dashed line are over- or underexpressed relative to the metagenomes, respectively. **(B)** Heatmap showing the relative abundance of ORFs in metatranscriptomes with highest similarity to Lokiarchaeon MAGs (n=982 ORFs), columns represent biological replicates. The dendrogram is a UPGMA based on the differences in gene expression between the hypoxic, interface, and sulfidic depths. This was statistically significant (ANOSIM: *P* = 0.002, R = 0.76). ORFs were classified into broad functional categories by comparison against the updated COG database^44^. Genes involved in the putative H_2_-dependent carbon fixation pathway are classified according to Sousa *et al*^4^. porABCD: 2-pyruvate:ferredoxin oxidoreductase, tungstopterin: Aldehyde ferredoxin oxidoreductase, HdrABC: heterodisulfide reductase, CDS/ACS: carbon monoxide dehydrogenase/acetyl-CoA synthase, mer: 5,10-methylene-H_4_-methanopterin reductase, ftr: formyl transferase, fwdB: formyl-methanofuran dehydrogenase, fdh: formate dehydrogenase, mtd: 5,10-methylene-H_4_-methanopterin dehydrogenase.

The incomplete nature of Asgard archaea genomes together with the lower representation of sequenced archaeal genomes compared to Bacteria^12^, make it likely that the high levels of archaeal gene expression seen here are actually an underestimate. Indeed, after updating our database with recently sequenced Lokiarchaeota MAGs^13,14^, the number of expressed ORFs in metatranscriptomes with highest similarity to Lokiarchaeota MAGs increased nearly two fold (from 522 to 982 ORFs). Lokiarchaeal MAGs comprise sequences from closely related strains and do not exhibit evidence of significant contamination^15^, thus we interpret expressed genes in our metatranscriptomes with highest similarity to Lokiarchaeon MAGs as being derived from the Lokiarchaeon group. However, we acknowledge that some genes in databases annotated as being present in Asgard archaea MAGs might not actually derive from asgard archaeal chromosomes^16^, but are present in environments where archaea occur and have been assigned to bins according to criteria that differ from study to study.

Expression of ORFs with similarity to those encoding the anaerobic H_2_-dependent WLP by Lokiarchaeon^5^ was observed predominantly in the sulfidic sediments (Fig 2b; CDS/ACS, mer, mtd, fwdB, ftr, fdh). Since Lokiarchaeon lacks the key gene methyl-CoM reductase for methanogenesis, the WLP likely functions as acetogenic (rather than methanogenic)^5^. Furthermore, ORFs with similarity to heterodisulfide reductases (HdrABC) involved in hydrogen based electron bifurcation^17^, and the coenzyme F420 NiFe hydrogenase (fhrB)^5^, were expressed primarily in the deepest interval sampled at 30 cmbsf (Fig. 2b). In methanogenic archaea, hydrogen dependent electron bifurcation uses the MvhADG-HdrABC system^17^ but we did not detect expression of MvhADG. Thus, it is possible that HdrABC based electron bifurcation in Lokiarchaeon proceeds in a different manner, for example *Desulfovibrio vulgaris* uses FlxABCD-HdrABC^18^. Autotrophs that fix CO_2_ via the WLP use pyruvate synthase (pyruvate:ferredoxin oxidoreductase, por) to generate pyruvate^5^ and Lokiarchaeon expression of this archaeal por was detected only in the interface and deepest (sulfidic) samples (Fig. 2b). These genes are typical of H_2_-dependent anaerobic autotrophs^5^, and their higher expression in the deepest (sulfidic) samples indicates a stimulation of hydrogen metabolism and the acetogenic WLP.

Since H_2_ production from fermentation^19^ is common in aquatic sediment environments, and high rates of sulfate reduction as an anaerobic terminal electron acceptor process have been measured in surface sediments throughout the Namibian shelf ^20^, it is reasonable to assume that in the anoxic (sulfidic) depths of the sediment core sampled in our study that H_2_ production and consumption is ongoing because this is a general feature of these environments^21^. H_2_ in anoxic sediments is produced by fermentation and tightly coupled and controlled by hydrogenotrophic bacteria and archaea using sulfate and bicarbonate as terminal electron acceptors^21,10^, but temporal disturbances (e.g., bioturbation, variability in overlying water mass oxidant composition, or variability in availability of carbon substrates) can lead to imbalances in H_2_ production and consumption rates. For instance, non-equilibrium H_2_ values >30 nmol/L have been observed in organic-rich continental slope sediments off Namibia^10^. Metabolic flexibility, such as possession of the WLP and capacity for utilization of organic substrates for fermentation may allow homoacetogenic Lokiarchaeon to take advantage of fluctuating H_2_ concentrations. Indeed, most probable number and ^14^C labeling experiments in similar sediments on the Chilean shelf suggested that variable redox cycling provided ideal conditions for homoacetogenic microorganisms^22^.

In addition to expression of the WLP, Lokiarchaeota expressed genes involved in the transport and metabolism of organic substrates, namely sugars and amino acids (Fig 2b). Glycolysis in Lokiarchaeon was evidenced by expression of ORFs with similarity to fructose-bisphosate aldolase and glyceraldehyde-3-phosphate dehydrogenase (Fig 2b: COG category G). Sugar fermentation by Lokiarchaeon is evidenced by the expression of ORFs with similarity to pyruvate-formate lyase that was expressed in the deepest samples (Fig 2b), which helps to regulate anaerobic glucose metabolism by catalyzing the reversible conversion of pyruvate and coenzyme-A into formate and acetyl-CoA^23^. Moreover, Lokiarchaeon expressed ORFs with similarity to an archaeal fructose-1,6-bisphosphate aldolase (Fig 2b: COG category G), which is induced during sugar fermentation in archaea and is responsible for sugar catabolism^24^.

Lokiarchaeon transcripts encoding amino acid and peptide transporters (Fig. 2b) further indicate mixotrophic utilization of detrital proteins. The expression of acryloyl-CoA reductase complex by Lokiarchaeon indicates that some of the amino acids are fermented, since this enzyme is used for L-alanine fermentation^25^. Amino acid fermentations would provide Lokiarchaeon with roughly one ATP per amino acid fermented^4^. This is consistent with amino acid fermentation^14,26,27^ and mixotrophy^28^ as a general metabolic features of anaerobic archaea in anoxic marine sediments. As these gene expression data suggested that Lokiarchaeota are anaerobic mixotrophic acetogens, or homoacetogens^8^, we tested their ability to use ^13^C-labeled organic matter and ^13^CO_2_ under anoxic conditions.

In the deepest (sulfidic) sample, where Lokiarchaeon gene expression was highest (Figs. 1, 2) we performed two sets of quantitative stable isotope probing (qSIP) experiments using ^13^C-labeled bicarbonate and ^13^C-labeled dEPS (see Methods) to identify autotrophic and heterotrophic activity, respectively. We used dEPS as a ^13^C-substrate for heterotrophic microbes because diatom derived organic matter substantially contributes to the total organic matter in Namibian shelf sediments^29^. The number of Asgard OTUs detected increased three fold at the end of the 10 day anoxic incubations compared to the *in situ* (frozen) samples, indicating that Asgard archaea were enriched for under the incubation conditions. For example, two Lokiarchaeon OTUs were detected in the frozen samples (Extended Data S3) whereas six Lokiarchaeon OTUs were detected at the end of the incubation (Fig. 3c). The extent of ^13^C enrichment in Asgard archaea populations from dEPS was high compared Bacteria, and nearly all Asgard populations were also labeled in the ^13^C-bicarbonate incubations suggesting mixotrophic activity (Fig. 3a). All Bathyarchaeota OTUs were significantly labeled in the ^13^C-bicarbonate incubations, and to a lesser extent in the ^13^C-labeled dEPS incubations compared to the Asgard populations (Fig. 3a). This experimental evidence supports previous SIP studies showing that Bathyarchaeota utilize organic matter marine sediments^30,31^, and the mixotrophic activity seen in our qSIP data supports genomic studies suggesting that Bathyarchaeota are organo-heterotrophic acetogens^32-34^. The high ^13^C-enrichment of the Lokiarchaeon and the Bathyarchaeota populations in the dEPS incubations, relative to the bacteria, also supports the significantly higher gene expression levels of these groups in the deepest sediment intervals (Fig. 2a).

**Figure 3:**
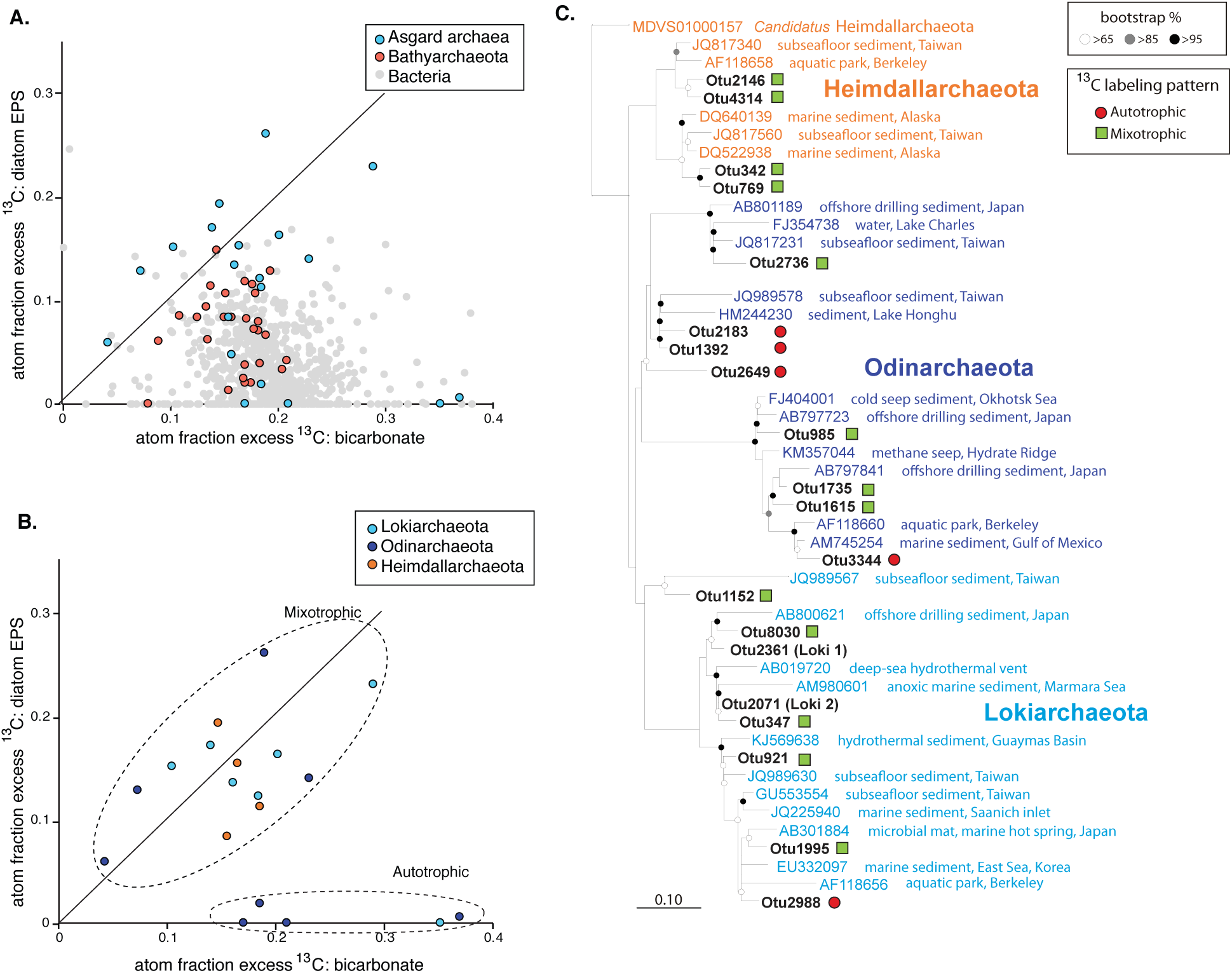
qSIP shows high activity and mixotrophy by Asgard archaea. (**A**) Atom fraction ^13^C with added ^13^C-bicarbonate and ^13^C-dEPS across all OTUs exhibiting significant labeling (90% CI not overlapping zero^45^) in at least one of the incubations. Each point represents the average atom fraction ^13^C values for separate OTU across three biological replicates, axes units represent percent of ^13^C-labeled carbon atoms in 16S rRNA genes per population in both incubations (0.25 indicates 25% of C atoms are ^13^C-labeled). (**B**) The same data, but displaying only Asgard archaea colored by candidate Phyla. The solid black lines indicate the expected relationship if organisms derived their carbon from bicarbonate and dEPS equally. The difference between the solid line and points falling above or below it is due to increased utilization of carbon from one of the other sources, and was used to define ^13^C-labeling patterns consistent with autotrophic and mixotrophic activity (dotted line circles). (**C**) Phylogenetic tree of 16S rRNA genes from Asgard OTUs (from panel B) that exhibited significant ^13^C-labeling patterns. Loki1 and Loki2 OTUs are populations of Lokiarchaeon recovered from the pristine frozen core (Extended Data Figure 3). A complete summary of the qSIP results for all taxa is presented in Extended Data Figure 4.

All OTUs affiliated with “*Candidatus* Lokiarchaeota” (n=6) were significantly enriched with ^13^C in the bicarbonate incubations (CI > 90%), with 5-35% total carbon atoms per Lokiarchaeon population being ^13^C -labeled (Fig. 3b). Five of the six Lokiarchaeon populations also exhibited a ^13^C-labeling pattern that was suggestive of mixotrophic activity, as they had similar labeling in both the bicarbonate and dEPS incubations (Fig. 3b). In comparison, an equal number of OTUs affiliated with the “*Candidatus* Odinarchaeota” exhibited ^13^C-labeling patterns consistent with mixotrophic and autotrophic activity, respectively (Fig. 3b). Three of these “*Candidatus* Odinarchaeota” OTUs exhibiting autotrophy formed a separate, bootstrap supported clade together with sequences from subseafloor and lake sediments (Fig. 3c). All detectable populations of “*Candidatus* Heimdallarchaeota” exhibited ^13^C-labeling patterns consistent with a mixotrophic metabolism (Fig. 3b, c). These results provide quantitative experimental support for genomic studies indicating that anaerobic mixotrophy is widespread in the Asgard archaea^6^ and show that some, particularly four populations affiliated with the “*Candidatus* Odinarchaeota” and one with Lokiarchaeon (Fig. 3b), may obtain the majority of their carbon from autotrophy. There is always a potential for cross-feeding in SIP experiments, whereby heterotrophs could feed off of the necromass of more active chemolithoautotrophs. Since the majority of OTUs in the ^13^CO_2_ incubations were identified as significantly incorporating the label (Fig S4), it is indeed possible that some cross feeding occurred over the 10 days in the ^13^CO_2_ incubation. However, it would be very difficult to explain the expression of the CO_2_ fixation WLP pathway by Lokiarchaeon (Fig. 2b), and the ^13^C-labeling pattern of Lokiarchaeon, if Lokiarchaeon was only getting labeled in the 13CO_2_ incubations by cross-feeding from more active autotrophic microbes.

The degree of ^13^C enrichment of Lokiarchaeon populations in the ^13^C-labeled dEPS incubations was higher than the bacterial average (Fig. 3a), providing quantitative evidence for their important role in the degradation of diatom necromass and dEPS in anoxic sediments. Clearly, the organic matter rich sediments^29,35^ on the Namibian shelf that we sampled represent an environment that promotes a high activity of Lokiarchaeon. Since polysaccharides are the main constituent of dEPS^36^ and we added this as a mixture of particulate and high molecular weight dissolved organic matter (HMW-DOM), Lokiarchaeon is apparently well suited to access polysaccharides from particulate and HMW-DOM. Mechanisms used by Lokiarchaeon to access this organic matter are evident in the metatranscriptomes via expression of ORFs with similarity to alpha-mannosidase (glycoside hydrolase family 38), pectate lyase (polysaccharide lyase family 9), cellobiose phosphorylase (glycoside hydrolase family 94) (Fig 2b: COG category G) that are all involved in the degradation of complex carbohydrates^37^. These enzymes help to degrade cellulose, pectin, and glycoproteins^37^ all of which are components of dEPS^38^. This heterotrophic activity of “*Candidatus* Lokiarchaeota” (previously referred to as the DSAG group) is consistent with a correlation of their abundance with organic matter in deeper subsurface sediment cores down to 3.5 meters below the seafloor^11^.

A coupling of H_2_-dependent CO_2_ reduction with H_2_-producing fermentations is common in homoacetogenic bacteria, that allows them to exploit otherwise-inaccessible organic substrates^8^. Homoacetogens couple the WLP with a variety of organic electron donors (e.g., sugars, ethanol, lactate), which is thought to be the most important ecological advantage of acetogens^8^. A good example is acetogenic sugar fermentation performed during homoacetate fermentation (“homoacetogenesis”), which provides the highest known ATP gain for glucose fermentations, more than 4.3 ATP/mol glucose in *A. woodii*^8^. This relatively high ATP yield could explain the high levels of anaerobic activity from “*Candidatus* Lokiarchaeota”, that are exhibiting a similar metabolism here, namely anaerobic utilization of carbohydrates, sugar fermentation, and the WLP. Our data indicate that Lokiarchaeota exhibit homoacetogenesis, coupling H_2_-producing fermentations with the H_2_-consuming WLP for CO_2_ fixation. This confirms earlier predictions for Lokiarchaeon metabolism based on metagenomics^33^ and adds to the growing evidence that homoacetogenesis is a widespread metabolism in marine sediments^34,39,40^.

The relatively high activity and anaerobic mixotrophy of Asgard archaea in organic rich, anoxic (sulfidic) Namibian shelf sediments, supports the genomic evidence that anaerobic mixotrophic metabolism is widespread in the candidate Asgard superphylum^4,6^. The Asgard archaea comprises the closest archaeal relatives of eukaryotes, and eukaryotes evolved during the Proterozoic eon when ocean anoxia was widespread^41,42^. H_2_ metabolism is furthermore hypothesized as a feature of the host cell that acquired the mitochondrion^43^. It is thus conceivable that H_2_-rich Proterozoic sediments under hypoxic waters with intense organic carbon flux to the seafloor (features of our sampled environment), would have potentially promoted the activity of LECA and represented an important habitat relevant to the emergence of the first eukaryotic cell.

## Acknowledgements

This work was supported by the Deutsche Forschungsgemeinschaft (DFG) through Project OR 417/4-1 (W.D.O), and the *F/S* Meteor Expedition M148/2 ‘EreBUS’. The authors thank the captain and crew of the *F/S* Meteor assistance during the oceanographic expedition, as well as S. Littmann, T. Wilkop, G. Klockgether and K. Imhoff who assisted in obtaining samples and providing chemical data. This work was performed in part through the Masters in Geobiology and Paleontology Program (MGAP) at LMU Munich. G.V.G-S was funded by the DFG project DI 842/6-1.

## Author contributions

W.D.O, conceived the idea for the study and wrote the paper. T.G.F. organized and led the expedition. A.V., P.R., O.K.C, G.V.G-S., V.M., and T. G. F. produced data. W.D.O., A.V., P.R., G.L., and O.K.C, analyzed data. T.G.F., G.V.G-S., and W.D.O acquired the samples.

## Competing interests

The authors declare no competing financial interests.

## Additional information

Supplementary Information includes Extended Data Fig S1 – S4 and Table S1. All sequence data is publicly accessible in NCBI through BioProject number PRJNA525353.

**Correspondence and requests for materials** should be addressed to W.D.O.

## Methods

### Sampling

A 30 cm long sediment core was obtained from a water depth of 125 m the Namibian continental shelf (18.0 S, 11.3 E) during *F/S* Meteor Expedition ‘EreBUS’ on July 10^th^, 2018. Sediments were sampled with a multi corer (diameter 10 cm), which yielded an intact sediment/water interface and the upper 30 cm of sediment. After retrieval, cores were moved immediately to a 4 °C cold room and sectioned every 2 cm within 24 hours. Sections were transferred immediately into sterile, DNA/RNA free 50 mL falcon tubes and then frozen immediately at -20 °C until DNA and RNA extractions. For the determination of dissolved sulfide and nitrate in the sediments, pore waters were anaerobically extracted at 12°C from multi-cores M148-206-6-8 and M148-206-5-8 using Rhizon samplers inserted at 3 to 5 cm intervals through pre-drilled holes along the side of the core-barrels. Dissolved oxygen in the water column was measured using two Clark-type Seabird Electronics (SBE 43) oxygen sensors on a wireline Seabird Electronics (USA) CTD SBE 911+ with rosette water sampler. Sensor values were calibrated by Winkler determinations of dissolved oxygen on discrete samples^43^.

### Pore water geochemistry

For sulfide analysis, 2 mL pore water aliquots were fixed with 0.5 mL of 5% (w/w) ZnCl_2_ (Fisher Scientific). Total dissolved sulfide, which included H_2_S_aq_ + HS^-^ + S^2+^ and the sulfidic component of polysulfides S_x_^2-^, was measured on board using the methylene blue method^46^ on a Shimadzu UV120 Spectrophotometer. Sulfide was measured from two separate 30 cm long cores collected via multi-coring. Nitrate was determined onboard with a QuAAtro39 autoanalyser (Seal Analytical) using the method based on Strickland and Parsons^47^.

### DNA and RNA extraction

DNA was isolated from 0.5 g sediment incubations aseptically in a sterile laminar flow hood using a sterile spatula, and extractions were carried out as described previously^48^. Purification of DNA extracts was carried out with the PowerClean Pro DNA Clean-up Kit (MO BIO Laboratories) and DNA was quantified with the Qubit dsDNA HS Assay kit (Thermo Fisher Scientific) according to manufacturer’s instructions. DNA was extracted using a previously described protocol. In addition, DNA from laboratory dust and extraction blanks was extracted and purified in order to identify and remove any contaminant sequences introduced during the laboratory processing of the samples.

Total RNA was extracted from 0.5 g of sediment using the FastRNA Pro Soil-Direct Kit (MP Biomedicals) following the manufacturer’s instructions with final elution of templates in 40 µL PCR water (Roche) as described previously^48^ with some modifications to maximize RNA yield and reduce DNA contamination. The first modification was that, after the supernatant was removed after first homogenization step, a second homogenization was performed with an additional 500 µL RNA Lysing Buffer. The tubes were centrifuged once again for 5 minutes at top speed, and the supernatant from the second homogenization was combined with that resulting from the first homogenization, continuing with the protocol from the manufacturer. Second, we added glycogen at a concentration of 1 µg/mL during the 30-minute isopropanol precipitation in order to maximize recovery of the RNA pellet. To reduce DNA contamination, we extracted all RNA samples in a HEPA-filtered laminar flow hood dedicated only for RNA work (no DNA allowed inside) that also contains dedicated RNA pipettors used exclusively inside the hood with RNA samples. All surfaces were treated with RNAse-Zap prior to extractions and exposed to UV light for 30 minutes before and after each extraction.

### qPCR and 16S rRNA gene sequencing

Bacterial 16S rRNA gene V4 hypervariable region was performed with primer pair 515F/806R using the qPCR protocol described previously^48^. In brief, qPCR was carried out in 20 µL solutions containing 10.4 µL Sso Advanced SYBR green PCR buffer (Bio-Rad, Hercules, CA, USA), 0.4 µL of 10 mM primer, 6.8 µL of nuclease-free water, and 2 µL of the DNA template. Three technical replicates were prepared with the epMotion 5070 robotic pipetting system (Eppendorf), with a technical variation of <5%. All reactions were performed with a two-step protocol in a CFX Connect real-time PCR system (Bio-Rad, Hercules, CA, USA), including an enzyme activation step at 95°C for 3 min, followed by 40 cycles of denaturation at 95°C for 15 s and then annealing at 55°C for 30 s. qPCR standards consisted of 10-fold dilution series of the genes of interest that were PCR amplified from the sample using the same primers. Prior to the creation of the dilution series, the amplified standard was gel extracted and quantified with a Qubit instrument. The reaction efficiencies in all qPCR assays were between 90% and 110%, with an r^2^ of 0.98. Gene copies were normalized to the wet weight of the sediment. Sequencing of the amplicons on the Illumina MiniSeq was performed as described previously^48^ through the LMU Munich GeoBio Center, which resulted in a total of 640,534 paired-end reads with an average of 23,000 reads per sample (SD: 3,000 reads) and USEARCH^49^ as described previously^48^. For 16S rRNA genes, taxonomic assignments were generated by QIIME, version 1.9.1^50^ using the implemented BLAST method against the SILVA rRNA gene database, release 132^51^. We removed all OTUs containing < 3 sequences and which had no BLASTn hit. All OTUs that were found in the dust and extraction blank (contamination) samples were also removed from the dataset. Reads passing this quality control were then normalized by percentage of total sequencing depth per sample. Two OTUs were preliminarily assigned to Lokiarchaea based on the taxonomy provided in the BLASTn searches against SILVA^51^, and confirmed using phylogenetics (Fig. S2a).

### Metatranscriptomics

DNAse treatment, synthesis of complementary DNA and library construction were obtained from 10 µL of RNA templates by processing the Trio RNA-Seq kit protocol (NuGEN Technologies). Libraries were quantified on an Agilent 2100 Bioanalyzer System, using the High Sensitivity DNA reagents and DNA chips (Agilent Genomics). The libraries constructed using specific (different) barcodes, pooled at 1 nM, and sequenced in two separate sequencing runs with a paired-end 300 mid output kit on the Illumina MiniSeq. A total of 40 million sequences were obtained after Illumina sequencing, which could be assembled *de novo* into 41,230 contigs. Quality control, *de novo* assembly, and ORFs searches were performed as described previously^52^. A total of 8,556 ORFs were found that were then searched for similarity using BLASTp against a database containing predicted proteins from all fungal, bacterial, and archaeal genomes and MAGs in the JGI and NCBI databases using DIAMOND^53^. This database included all Asgard and Lokiarchaetoa MAGs from recent studies^13,14^, corresponding to 61,913 predicted proteins from Lokiarchaeota (as of July 26^th^, 2019). It also contained all ORFs assigned as Lokiarchaeota from the metagenomes prepared from the same samples, as well as all ORFs from the >700 transcriptomes of microbial eukaryotes from the MMETS project^54^. Cutoff for assigning hits to specific taxa were a minimum bit score of 50, minimum amino acid similarity of 30, and an alignment length of 50 residues. Extraction blanks were also sequenced alongside the environmental samples to identify contamination, and ORFs from contaminant taxa. Contamination in the metatranscriptomes were primarily diatoms (“lab weeds”), cyanobacteria, *Streptococcus, Acinetobacter, Staphylococcus, Rhizobium, Ralstonia,* and *Burkholderia.* All ORFs that were shared between contaminant samples and the metatranscriptomes were removed prior to analysis. Incorporation of protist transcriptomes^54^ greatly reduced the amount of laboratory contamination from eukaryotic algae such as diatoms (“lab weeds”) introduced during the library prep. All metatranscriptomes had <10% ORFs from contaminating taxa.

### Metagenomics

DNA was extracted according to the protocol described previously, which recovers ca. 10 times more DNA per gram sediment extracted compared to conventional protocols^55^. The protocol yields more DNA because it involves a phosphate buffer that prevents DNA chelation to clay mineral surfaces, employs multiple freeze thaw steps, and concentrates the DNA with Amicon filters as opposed to precipitating with ethanol or isopropanol. This yielded microgram quantities of DNA from the core top, 12 cm, and 28 cm samples (1 g sediment was used for the extraction) and allowed us to create Illumina ready libraries from microgram quantities of extracted DNA without prior amplification. Illumina libraries were prepared, quality checked, and sequenced as described previously^55^. De novo assembly of raw reads passing quality control, ORF finding, and annotation were performed using the same pipeline applied previously for sediment metagenome datasets^55^ and the metatranscriptomes in this study (see above section). This database included all publicly available Asgards MAGs to date in the NCBI Protein database (corresponding to 61,913 predicted proteins from Lokiarchaeota as of July 26^th^, 2019). Extraction blanks were also sequenced alongside the environmental samples to identify contamination, and ORFs from contaminant taxa were removed prior to analysis. All metagenomes had <1% ORFs from contaminating taxa.

As an additional method for assessing the relative abundance of Lokiarchaeota in the metagenomes, raw reads from the were mapped against the assembled MAG of Lokiarchaeum sp. GC14_75, GenBank contig references no. JYIM010000001.1 to 505.1 (Spang *et al*., 2015)^2^ using Geneious v. 8.1.9. For this genome mapping, we ran 100 iterations at medium sensitivity, which entailed the following parameters: minimum mapping quality: 25%, minimum coverage: 5 reads, strict threshold: 50%. An average of 2 % (+/- 0.6 %) reads mapped to the assembled Lokiarchaeota MAG^2^, which covered the entire genome (504/504 contigs recovered at all depths). This was comparable to the relative abundance of *de novo* assembled metagenome contigs encoding ORFs with highest similarity to Lokiarchaeota MAGs, which was 1.8 % (+/- 0.7%). Thus, both methods (genome recruitment vs. de novo assembly) indicate the same relative abundance for Lokiarchaeota at ca. 2% of the total community.

### Normalizing gene expression

Group-specific gene expression levels were calculated as follows. The relative abundance of ORFs (defined as percent of total ORFs) with highest similarity to a particular group were normalized against the same relative abundance in the metagenome from the same sample. For this normalization, we only considered ORFs that had similarity to a database entry at relatively high confidence (see above). We chose to focus on total ORFs detected, as opposed to number of reads mapping per kilobase per ORF (e.g., RPKM), in order to reduce potential bias from small numbers of ‘housekeeping’ genes with potentially higher expression levels. Thus, by considering total number of ORFs expressed (as opposed RPKM values), the normalization displays the percentage of the pan-genome that is expressed as a function of group abundance. The normalized values are thus placed on a scale of either greater or less than 1, corresponding to either over- or underexpression, respectively. In other words, if the relative abundance of expressed ORFs from a particular group is higher in the metatranscriptome relative to the metagenome, then the value will be above 1. This calculation represents a group average in activity a semi-quantitative approach to compare the average level of transcriptional activity between Phylum level groups in the communities sampled.

### Experimental set up for ^13^C incubations

Sediments from a depth of 28 cmbsf were incubated in triplicate for 10 days in the dark with either 2 mM 99% ^13^C-labeled or unlabeled (control) sodium bicarbonate (NaHCO_3_; Sigma-Aldrich, St. Louis, MO, USA) at 10 °C. A second set of incubations from the same sediment depth were set up with either ^13^C-labeled or unlabeled (control) diatom necromass / EPS, produced from a culture of the diatom *Chaetocerous socialis* (Norwegian Culture Collection strain K1676). *C. socialis* cells were grown in 250 mL sterile polystyrene culture flasks (VWR International) with L1 growth medium^56^ at 22 °C for seven days exposed to the natural light dark cycle (flasks were placed in an east-facing window). One set of cultures was grown with 2 mM 99% ^13^C-labeled sodium bicarbonate and another set was grown with unlabeled sodium bicarbonate. After seven days the cultures were turbid, as evidenced by mucosal light brown flocculant (EPS and colonies of diatom cells) and the cultures at that point were concentrated in 50 KDa Amicon filters. Thus, the resulting cell culture concentrates consisted primarily of particulate and high-molecular-weight dissolved organic matter. GC-IRMS was used to determine that the atom percent ^13^C enrichment of the organic matter was >50%. For the incubations, flasks received either the unlabeled or ^13^C-labeled organic matter at a final concentration of 200 µg per gram. As the total organic carbon content of sediments at a several nearby locations (18.3 S, 11.5 E) was determined previously to be between 0.5 and 2.3%^35^, we added labeled organic matter at a concentration roughly equal to 1-3% of the *in situ* concentration. Since sinking diatom biomass is a major contributor to the organic carbon content of Namibian shelf sediments^29^, the ^13^C-labeled organic matter representing a mixture of dead diatom cells and their EPS serves as an appropriate proxy for tracking activity of heterotrophic microbes in the sediments.

### Quantitative stable isotope probing

Sediment was added to 20 mL glass flasks leaving no headspace (ca. 20 g sediment) that were crimp sealed using grey butyl rubber stoppers. Anoxic conditions were confirmed by monitoring O_2_ concentrations non-invasively as described previously^48^ using a Fibox fiber optic O_2_ sensor. After 10 days the flasks were frozen until processing for SIP. DNA from the samples was extracted as described above and prepared for density gradient centrifugation according to the quantitative stable isotope probing protocol as described previously^45^. In brief, density gradient centrifugations were carried out in a TLN-100 Optima MAX-TL ultracentrifuge (Beckman Coulter, Brea, CA, USA) near-vertical rotor at 18 °C for 72 h at 165,000 × *g*. 50 µL of DNA spanning from 0.5 µg to 1.5 µg which was within the range of proposed values^57^ was added to a solution of cesium chloride (CsCl) and gradient buffer (0.1 M Tris, 0.1 M KCl and 1 mM EDTA) in order to achieve a starting density of 1.70 g mL^-1^ in a 3.3 mL polyallomer OptiSeal tubes (Beckman Coulter, Brea, CA, USA). After ultracentrifugation, the density gradients were fractionated into 15 equal fractions of 200 µL from the bottom of polyallomer OptiSeal tubes by using a syringe pump and fraction recovery system (Beckman Coulter, Brea, CA, USA). The density of these fractions was measured with an AR200 digital refractometer (Reichert Analytical Instruments, Depew, NY, USA). DNA was precipitated from the fractions using 2 volumes of polyethylene glycol with 2 µL (10 mg mL^-1^) glycogen and precipitated overnight at room temperature. DNA was pelleted by centrifugation (13,000 × *g*; 40 min), washed with 70% ethanol, and resuspended with 30 µL molecular-grade (DEPC-treated) water. DNA was quantified fluorometrically using a Qubit. The observed excess atom ^13^C-enrichment fraction (EAF) was calculated for each taxon according to a previously described study^45^ using a qSIP workflow embedded in the HTS-SIP R package^58^. Weighted average densities were calculated for each taxon’s DNA in the control (^12^C added) and in the experimental incubation (^13^C added) as described Hungate *et al.* ^59^ to estimate the atom fraction excess of ^13^C for each OTU. To calculate the bootstrap confidence intervals (CI) for significant isotopic incorporation, bootstrap replicates (n= 1000) were run with the HTS-SIP R package ^58^; an OTU was considered to be ^13^C labeled if the 90% CI was above the 0% EAF cutoff ^59^.

